# LAB–QA2GO: A free, easy-to-use toolbox for the quality assessment of magnetic resonance imaging data

**DOI:** 10.1101/546564

**Authors:** Christoph Vogelbacher, Miriam H. A. Bopp, Verena Schuster, Peer Herholz, Andreas Jansen, Jens Sommer

## Abstract

Image characteristics of magnetic resonance imaging (MRI) data (e.g. signal-to-noise ratio, SNR) may change over the course of a study. To monitor these changes a quality assurance (QA) protocol is necessary. QA can be realized both by performing regular phantom measurements and by controlling the human MRI datasets (e.g. noise detection in structural or movement parameters in functional datasets). Several QA tools for the assessment of MRI data quality have been developed. Many of them are freely available. This allows in principle the flexible set-up of a QA protocol specifically adapted to the aims of one’s own study.

However, setup and maintenance of these tools bind time, in particular since the installation and operation often require a fair amount of technical knowledge. In this article we present a light-weighted virtual machine, named *LAB-QA2GO*, which provides scripts for fully automated QA analyses of phantom and human datasets. This virtual machine is ready for analysis by starting it the first time. With minimal configuration in the guided web-interface the first analysis can start within 10 minutes, while adapting to local phantoms and needs is easily possible. The usability and scope of *LAB–QA2GO* is illustrated using a data set from the QA protocol of our lab. With *LAB–QA2GO* we hope to provide an easy-to-use toolbox that is able to calculate QA statistics without high effort.

## 1 Introduction

Over the last 30 years, magnetic resonance imaging (MRI) has become an important tool both in clinical diagnostics and in basic neuroscience research. Although modern MRI scanners generally provide data with high quality (i.e. high signal-to-noise ratio, good image homogeneity, high image contrast and minimal ghosting), image characteristics will inevitably change over the course of a study. They also differ between MRI scanners, making multicenter imaging studies particularly challenging (Vogelbacher et al., 2018). For longitudinal MRI studies stable scanner performance is required not only over days and weeks, but over years, for instance to differentiate between signal changes that are associated with the time course of a disease and those caused by alterations in the MRI scanner environment. Therefore, a comprehensive quality assurance (QA) protocol has to be implemented that monitors and possibly corrects scanner performance, defines benchmark characteristics and documents changes in scanner hardware and software (Glover et al., 2012). Furthermore, early-warning systems have to be established that indicate potential scanner malfunctions.

The central idea of a QA protocol for MRI data is the regular assessment of image characteristics of a MRI phantom. Since the phantom delivers more stable data than living beings, it can be used to disentangle instrumental drifts from biological variations and pathological changes. Phantom data can be used to assess, for instance geometric accuracy, contrast resolution, ghosting level, and spatial uniformity. Frequent and regular assessments of these values are needed to detect gradual and acute degradation of scanner performance. Many QA protocols additionally complement the assessment of phantom data with the analysis of human MRI datasets. For functional imaging studies, in which functional signal changes are typically just a small fraction (~1-5 %) of the raw signal intensity (Friedman and Glover, 2006), in particular the assessment of the temporal stability of the acquired time series is important, both within a session and between repeated measurements. The documented adherence to QA protocols has therefore become a key benchmark to evaluate the quality, impact and relevance of a study (Van Horn and Toga, 2009).

Different QA protocols for MRI data are described in the literature, mostly in the context of large-scale multicenter studies (for an overview, see Glover et al. (2012) and Van Horn and Toga (2009)). Depending on the specific questions and goals of a study, these protocols typically focused either on the quality assessment for structural (e.g. Gunter et al. (2009)) or functional MRI data (e.g. Friedman and Glover (2006)). QA protocols were also developed for more specialized study designs, for instance in multimodal settings as the combined acquisition of MRI with EEG (Ihalainen et al., 2015) or PET data (Kolb et al., 2012). Diverse MRI phantoms are used in these protocols, e.g. the phantom of the American College of Radiology (ACR) (ACR, 2005), the Eurospin test objects (Firbank et al., 2000) or gel phantoms proposed by the Functional Bioinformatics Research Network (FBIRN)-Consortium (Friedman and Glover, 2006). These phantoms were designed for specific purposes. Whereas for instance the ACR phantom is well suited for testing the system performance of a MRI scanner, the FBIRN phantom was primarily developed for fMRI studies.

A wide array of QA algorithms is used to describe MR image characteristics, for instance the so-called “Glover parameters” applied in the FBIRN consortium (Friedman and Glover, 2006) (for an overview see Glover et al. (2012) and Vogelbacher et al. (2018)). Many algorithms are freely available (see e.g. C-MIND(Lee et al.), CRNL (Chris Rorden’s Neuropsychology Lab (CRNL) | MRI Imaging Quality Assurance Methods); ARTRepair (Mazaika et al., 2009); C-PAC (Cameron et al., 2013)). This allows in principle the flexible set-up of a QA protocol specifically adapted to the aims of one’s own study. The installation of these routines, however, is often not straight-forward. It typically requires a fair level of technical experience, e.g. to install additional image processing software packages or to handle the dependence of the QA tools on specific software versions or hardware requirements^1^. We therefore developed an easy-to-use QA tool which provides on the one hand a fully automated QA pipeline for MRI data, but is on the other hand easy to integrate on most imaging systems and does not require particular hardware specifications. In this article we present the main features of our QA tool, named *LAB–QA2GO.* In the following, we give more information on the technical implementation of the *LAB–QA2GO* tool (section 2), present a possible application scenario (“center specific QA”) (section 3) and conclude with an overall discussion (section 4).

## 2 Technical implementation of *LAB-QA2GO*

In this section, we describe the technical background of *LAB-QA2GO*, outline different QA pipelines and describe the practical implementation of the QA analysis. These technical details are included as part of a manual in a MediaWiki (version: 1.29.0, https://www.mediawiki.org/wiki/MediaWiki/de) as part of the virtual machine. The MediaWiki could also serve for the documentation of the laboratory and/or study.

### 2.1 Technical background

*LAB–QA2GO* is a virtual machine (VM)^2^. Due to the virtualization, the tool is already fully configured and easy to integrate in most hardware environments. All functions for running a QA analysis are installed and immediately ready-for-use. Also all additionally required software packages (e.g. FSL) are preinstalled and preconfigured. Only few configuration steps have to be performed to adapt the QA pipeline to own data. Additionally, we developed a user-friendly web interface to make the software easily accessible for inexperienced users. The VM can either be integrated into the local network environment to use automatization steps or it can be run as a stand-alone VM. By using the stand-alone approach, the MRI data has to be transferred manually to the *LAB–QA2GO* tool. The results of the analysis are presented on the integrated web based platform (Fig. 1). The user can easily check the results from every workstation (if the network approach is chosen).

**Fig. 1:**
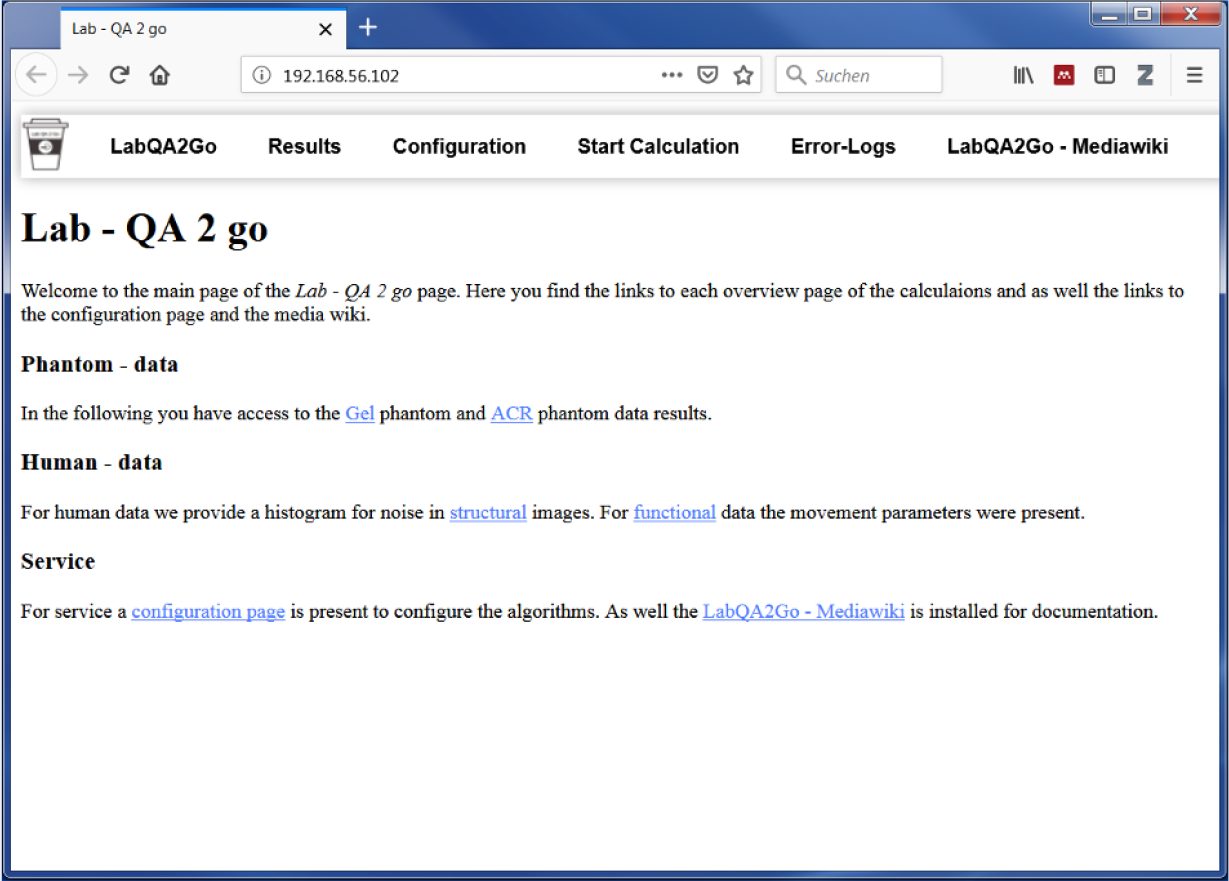
The web based graphical user interface of the LAB–QA2GO tool presented in a web browser.

We choose NeuroDebian (http://neuro.debian.net, version: 8.0) as operating system for the VM, as it provides a large collection of neuroscience software packages (e.g. Octave, mricron) and has a good standing in the neuroscience community. To keep the machine small, i.e. the space required for the virtual drive, we included only packages necessary for the QA routines in the initial setup and decided to use only open source software. But users are free to add packages according to their needs. To avoid license fees, we opted to use only open source software. The full installation documentation can be found in the MediaWiki of the tool.

For providing a web based user-friendly interface, presenting the results of the QA pipelines and receiving the data, the light-weight ligthhtpd webserver (version: 1.4.35, https://www.lighttpd.net) is used. The web based interface can be accessed by any web browser (e.g. the web browser of the host or the guest system) using the IP address of the *LAB–QA2GO* tool. This webserver needs little hard disk space and all required features can easily be integrated. The downscaled Picture Archiving and Communication System (PACS) tool Conquest (version: 1.4.17d, https://ingenium.home.xs4all.nl/dicom.html) is used to receive and store the Digital Imaging and Communications in Medicine (DICOM) files. Furthermore, we installed PHP (version: 5.6.29-0, http://php.net) to realize the user interface interaction. Python (version: 2.7.9, https://www.python.org/) scripts were used to for the general schedule, to move the data into the given folder structure, to start the data specific QA scripts, collect the results and write the results into HTML files. The received DICOM files were transferred into the Neuroimaging Informatics Technology Initiative (NIfTI) format using the dcm2nii tool (version: 4AUGUST2014 (Debian) https://www.nitrc.org/projects/dcm2nii/). To extract the DICOM header information, the tool dicom2 (version: 1.9n, http://www.barre.nom.fr/medical/dicom2/) is used, which converts the DICOM header into an easy accessible and readable text file.

The QA routines were originally implemented in Matlab (www.mathworks.com) (Hellerbach, 2013; Vogelbacher et al., 2018) and got adapted to GNU Octave (version: 3.8.2, www.octave.de) for *LAB–QA2GO.* The NeuroImaging Analysis Kit (NIAK) (version: boss-0.1.3.0, https://www.nitrc.org/projects/niak/) was used for handling the NIfTI files and graphs were plotted using matplotlib (version: 1.4.2, https://matplotlib.org/index.html), a plotting library for python.

Finally, to process human MRI data we used the image processing tools of FMRIB Software Library (FSL, version: 5.0.9, https://fsl.fmrib.ox.ac.uk/fsl/fslwiki). Motion Correction FMRIB’s Linear Image Registration Tool (MCFLIRT) was used to compute movement parameters of fMRI data and Brain Extraction Tool (BET) to get a binary brain mask.

### 2.2 QA pipelines for phantom and for human MRI data

Although the main focus of the QA routines was on phantom datasets, we added a pipeline for human datasets (raw DICOM data from the MR scanner). To specify which analysis should be started, *LAB–QA2GO* uses unique identifiers to run either the human or the phantom QA pipeline.

<u>For phantom data analysis</u>, *LAB–QA2GO* runs an automated QA analysis on data of an ACR phantom and a gel phantom (for an overview see Glover et al.(2012)). Additional analyses, however, can be easily integrated in the VM. For the analysis of ACR phantom data, we used the standard ACR protocol (ACR, 2005). For the analysis of gel phantom data, we used statistics previously described by Friedman et al. (2006) (the so-called “Glover parameters”), Simmons et al. (1999) and Stöcker et al. (2005).

These statistics assess e.g. the signal-to-noise ratio, the uniformity of an image or the temporal fluctuation. Detailed information on these statistics can be found elsewhere (Vogelbacher et al., 2018). In Fig. 2 the calculation of the signal-to-noise-ratio (SNR) based on the “Glover parameters” is shown exemplarily.

**Fig. 2:**
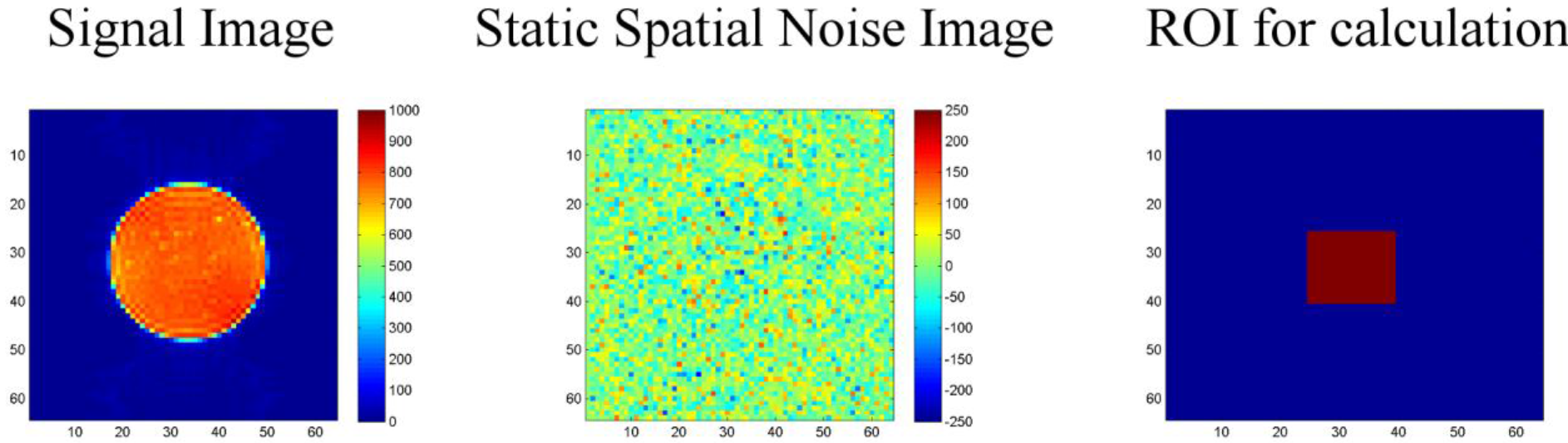
Calculation of the signal-to-noise-ratio (SNR) for the gel phantom. First, a “signal image” is calculated as the voxel-wise average of the center slice of the gel phantom (slice-of-interest, SOI) across the time series (left). Second, a “static spatial noise image” is calculated as the voxel-wise difference of the sum of all odd images and the sum of all even images in the SOI (middle). Third, the SNR is defined as the quotient of the average intensity of the mean signal image in a region of interest (ROI, 15 x 15 voxel), located in the center of the phantom of the SOI (right), and the standard deviation of the static spatial noise within the same ROI.

<u>For human data analysis</u>, we use movement parameters from fMRI and noise level from structural MRI as easily interpretable QA parameters (Fig. 3). The head movement parameters (translation and rotation) are calculated using FSL MCFLIRT and FSL FSLINFO with default settings, i.e. motion parameters relative to the middle image of the time series. Each parameter (pitch, roll, yaw, movement in x, y and z direction) is plotted for each time-point in a graph (Fig. 3 top). Additionally, a histogram is generated of the step width between two consecutive time points to detect large movements between two time points (Fig. 3 middle). For structural MRI data, a brain mask is calculated by FSL’S BET (using the default values) first. Subsequently, the basal noise of the image background (i.e. the area around the head) is detected (Fig. 3 bottom). The calculated mask is saved to create images for the report. Also a basal SNR (bSNR) value is calculated by the quotient of the mean intensity in the brain mask and the standard deviation of the background signal. Each value is presented individually in the report to easily see by which parameter the SNR value was influenced. These two methods should give the user an overview of existing noise in the image. Both methods can be independently activated or deactivated by the user to individually run the QA routines.

**Fig. 3:**
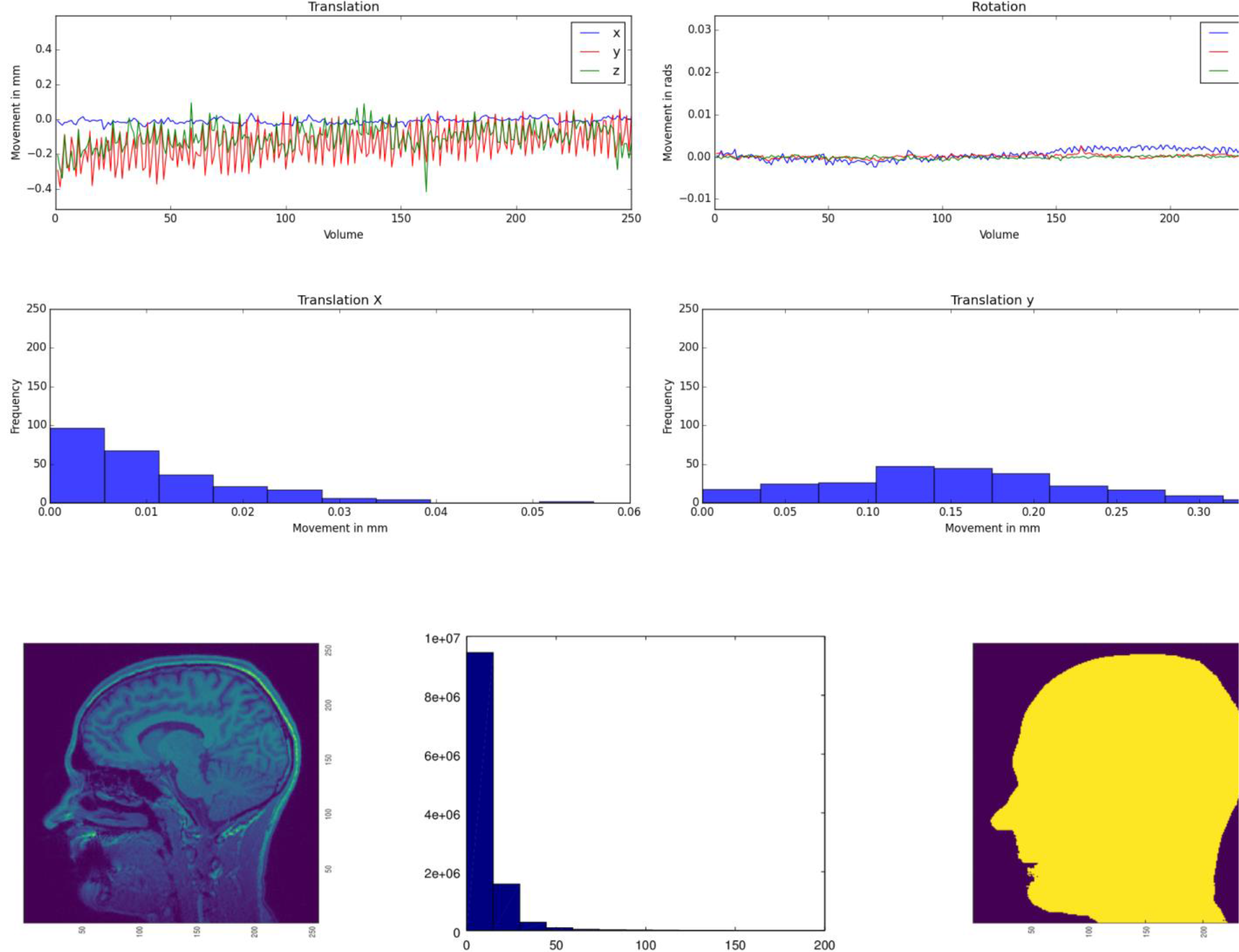
Top: Head movement parameters (translation and rotation) for functional MRI data. Center: Histogram of the translation in x and y direction. Middle: The movement parameters are transferred into a histogram, illustrating the amount of movement between two consecutive time points (exemplarily shown for the x and y translation parameters). Bottom: Noise calculation algorithm for structural MRI data. First, a region of interest (ROI) is defined in the corner of the three-dimensional image. Second, the mean of this ROI aggregated by an initial user defined threshold multiplier is used to mask the head in a first step (idea: use the intensity of the scalp to get a first approximation). Third, for every axial and sagittal slice the edges of the scalp were detected by using a differentiation algorithm between two images to create a binary mask of the head. Fourth, this binary mask is multiplied with the original image to get the background of the head image. Fifth, for this background a histogram of the containing intensity values is generated.

### 2.3 Practical implementation of QA analyses in *LAB-QA2GO*

The *LAB-QA2GO* pipelines (phantom data, human data) are preconfigured, but require unique identifiers as part of the dicom field ‘patient name’ to distinguish between data sets, i.e. which pipeline should be used for the analysis of the specific data set. Predefined are ‘ Phantom’, ‘ACR’ and ‘GEL’ in the field ‘patient name’, but can be adopted to the local needs. These unique identifiers have to be inserted into the configuration page (a web based form) on the VM (Fig. 4). The algorithm checks for the field “patient name” of the DICOM header so that the unique identifier has to be part of the “patient name” and has to be set during the registration of the patient at the MR scanner.

**Fig. 4:**
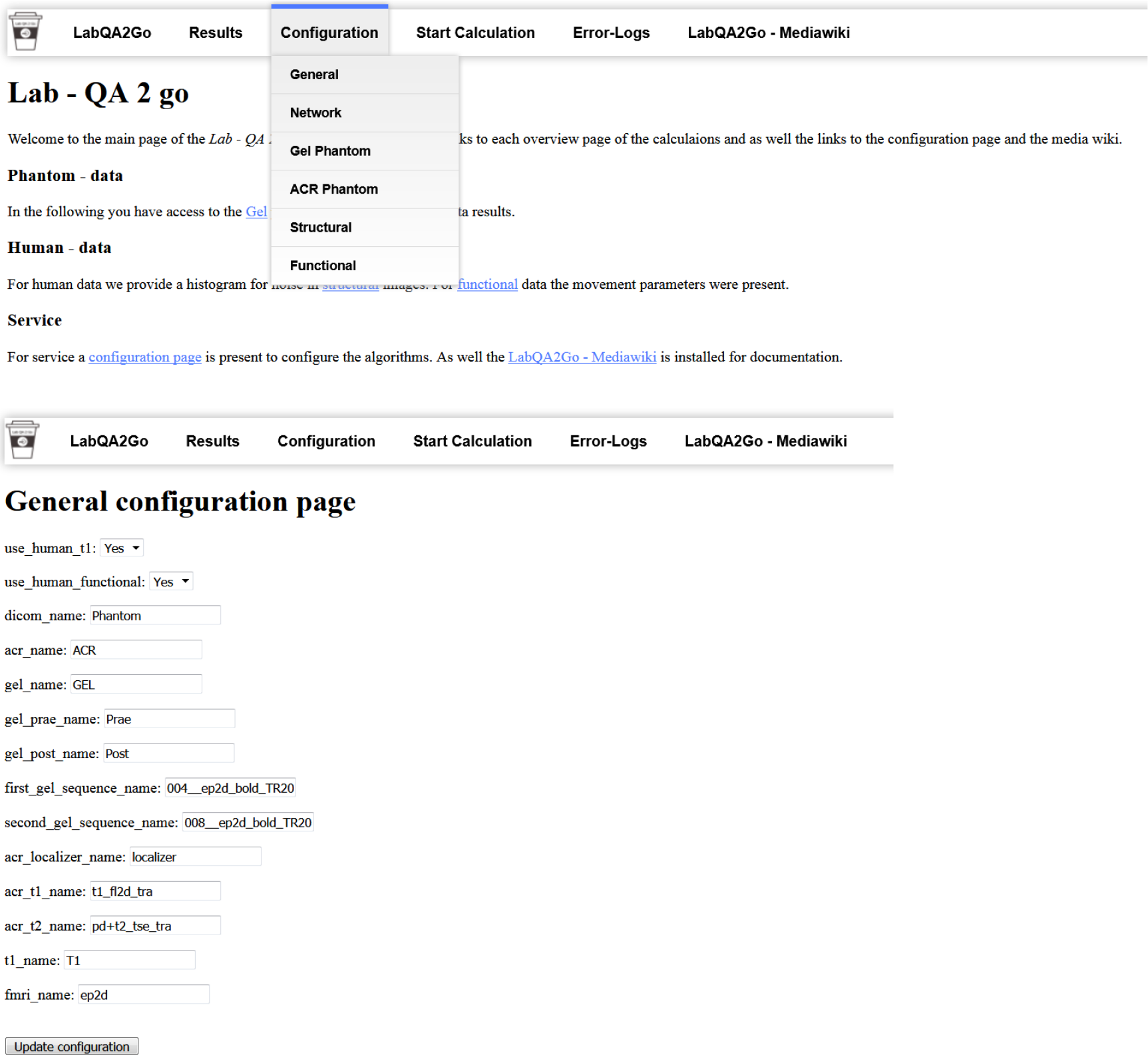
The general configuration page used to set the unique identifier and to activate or deactivate the human dataset QA pipeline.

The MRI data are integrated into the VM either by sending them (“dicom send”, network configuration) or providing them manually (directory browsing, stand-alone configuration). Using the network configuration, the user has to integrate the IP address of the VM as a DICOM receiver in the PACS first. *LAB–QA2GO* runs the Conquest tool as receiving process to receive the data from the local setup, i.e. either the MRI camera, the PACS, etc. and stores them in the VM. Using the stand-alone configuration, the user has to copy the data manually to the VM. This can be done using e.g. a USB-Stick or a shared folder with the host system (provided by the virtualization software). In the stand-alone configuration, the VM can handle both DICOM and NIfTI format data. The user has to specify the path to the data in the provided web interface and then just press start. If the data is present as DICOM files, then the DICOM data send process is started to transfer the DICOM files to the conquest tool, to run the same routine as described above. If the data is present in NIfTI format, the data is copied into the temporal folder and the same routine is started without converting the data.

After the data is available in the *LAB–QA2GO* tool, the main script for analysis is either started automatically at a chosen time point or can be started manually by pressing a button in the web interface. The data processing is visualized in Fig. 5. First, the data is copied into a temporal folder. Data processing is performed on NIfTI formatted data. If the data is in DICOM format, it will be converted into NIfTI format using the dcm2nii tool. Second, the names of the NIfTI files are compared to the predefined unique identifiers. If the name of the NIfTI data partly matches with a predefined identifier, then the corresponding QA routine is started (e.g. gel phantom analysis; see section 2.2).

**Fig. 5:**
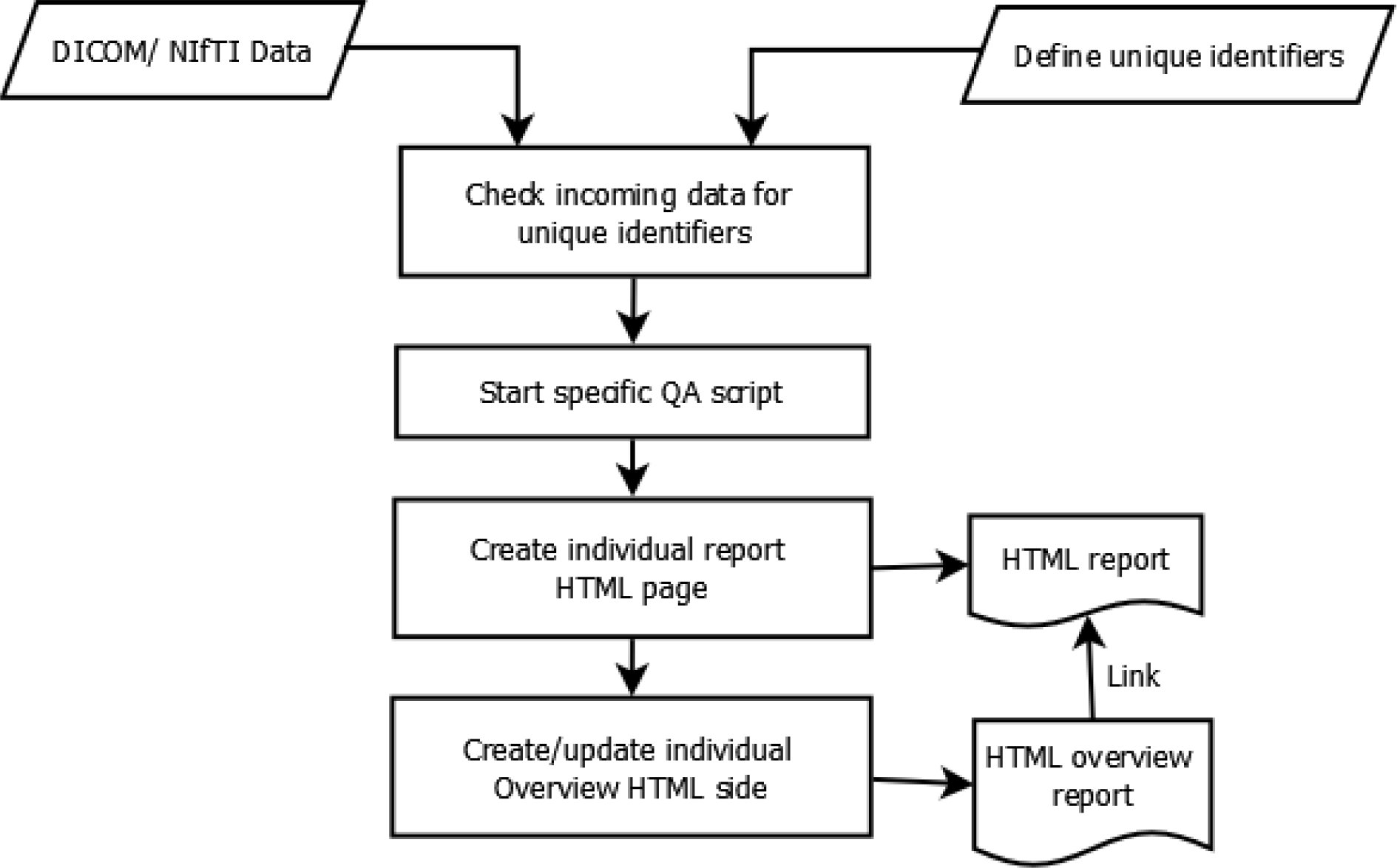
Data flow diagram of the QA analysis

Third, after each calculation step, a HTML file for the analyzed dataset is generated. In this file, the results of the analysis are presented (e.g. the movement graphs for functional human datasets). In Fig. 6, we show an exemplary file for the analysis of gel phantom data. Furthermore, an overview page for each analysis type is generated or updated. On this overview page, the calculated parameters of all measurements of one data type are presented as a graph. An individual acceptance range can be defined using the configuration page, which is visible in the graph. Additionally, all individual measurement result pages are linked at the bottom of the page for a detailed overview. Outliers (defined by the acceptance range) are highlighted to detect them easily.

**Fig. 6:**
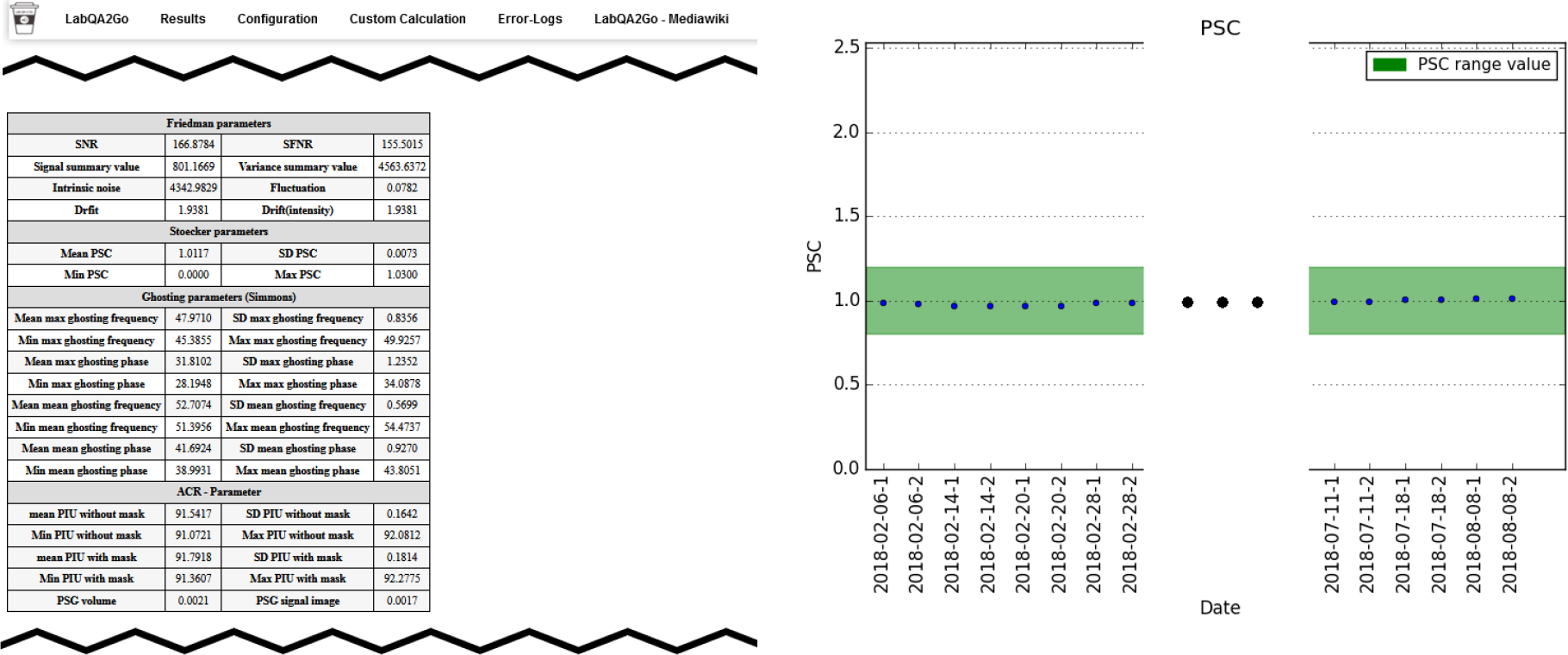
Excerpt from the results page summarizing the results of a QA analysis of a gel phantom. Left: Summary of all QA statistics measured at a specific time point. Right: Overview of the percent signal change (PSC) of the MRI signal over a period of 6 months. This graphic shows stable MRI scanner performance.

## 3 Application scenario: Quality assurance of an MRI scanner

There are many possible application scenarios for the *LAB–QA2GO* tool. It can be used, for instance, to assess the quality of MRI data sets acquired in specific neuroimaging studies (e.g. Frässle et al.(2016)) or to compare MRI scanners in multicenter imaging studies (e.g. Vogelbacher et al. (2018)). In this section we will describe another application scenario in which the *LAB–QA2GO* tool is used to assess the long-term performance of one MRI scanner (“center-specific QA”). We will illustrate this scenario using data from our MRI lab at the University of Marburg. The aim of this QA is not to assess the quality of MRI data collected in a specific study, but to provide continuously information on the stability of the MRI scanner across studies.

### Center-specific QA protocol

The assessment of MRI scanner stability at our MRI lab is based on regular measurements of both the ACR phantom and a gel phantom. The phantoms are measured at fix time points. The ACR phantom is measured every Monday and Friday, the gel phantom each Wednesday. All measurements are performed at 8 a.m., as first measurement of the day. For calculating the QA statistics, the *LAB–QA2GO* tool is used in the network configuration. As unique identifiers (see section 2), we determined that all phantom measurements must contain the keywords “phantom” and either “GEL” or “ACR” in the “patient name”. If these keywords are detected by *LAB–QA2GO*, the processing pipelines for the gel phantom analysis and the ACR phantom analysis, respectively, are started automatically. In the following, we describe the phantoms and the MRI protocol in detail. We also present examples how the QA protocol can be used to assess the stability of the MRI scanner.

#### Gel phantom

The gel phantom is a 23.5 cm long and 11.1 cm-diameter cylindrical plastic vessel (Rotilabo, Carl Roth GmbH + Co. KG, Karlsruhe, Germany) filled with a mixture of 62.5 g agar and 2000 ml distilled water. In contrast to widely used water filled phantoms, agar phantoms are more suitable for fMRI studies. On the one hand, T2 values and magnetization transfer characteristics are more similar to brain tissue (Hellerbach, 2013). Furthermore, gel phantoms are less vulnerable to scanner vibrations and thus avoid a long settling time prior to data acquisition (Friedman and Glover, 2006). For the gel phantom, we chose MR sequences that allowed to assess the temporal stability of the MRI data. This stability is in particular important for fMRI studies in which MRI scanners are typically operated close to their load limits. The MRI acquisition protocol consists of a localizer, a structural T1-weighted sequence, a T2*-weighted echo planar imaging (EPI) sequence, a diffusion tensor imaging (DTI) sequence, another fast T2*-weighted EPI sequence and, finally, the same T2*-weighted EPI sequence as at the beginning. The comparison of the quality of the first and the last EPI sequence allows in particular to assess the impact of a highly stressed MRI scanner on the imaging data. The MRI parameters of all sequences are listed in Table 1.

**Table 1:** MRI parameters for the gel phantom measurements

| **Sequence**<br>**(position in QA**<br>**protocol)** | **Localizer (1)** | **T1(2)** | **bold**<br>**sensitive**<br>**EPI (3, 6)** | **diffusion**<br>**sensitive**<br>**EPI (4)** | **bold**<br>**sensitive**<br>**EPI (5)** |
| --- | --- | --- | --- | --- | --- |
| Repetition time (TR) | 8.6 ms | 1900 ms | 2000 ms | 3900 ms | 177 ms |
| Echo time (TE) | 4 ms | 2.26 ms | 30 ms | 90 ms | 30 ms |
| Field of View (FoV) | 250 mm | 256 mm | 210 mm | 256 mm | 210 mm |
| Matrix size | 256*256 | 256 *256 | 64*64 | 128*128 | 64*64 |
| Slice thickness | 7.0 mm | 1.0 mm | 3.0 mm | 2.0 mm | 3.0 mm |
| Distance factor |  | 50% | 20% | 0% | 20% |
| Flip angle | 20 ° | 9° | 70° |  | 70° |
| Phase encoding direction | anterior >> posterior ,<br>anterior >> posterior, right >> left | anterior >> posterior | anterior >> posterior | anterior >> posterior | anterior >> posterior |
| Bandwidth (Hz/Px) | 320 | 200 | 2894 | 1502 | 2894 |
| Acquisition order (series) | Sequential (interleaved) | Single shot (ascending) | Interleaved (interleaved) | Interleaved (interleaved) | Interleaved (interleaved) |
| Number of slices | 2 in each direction | 176 | 34 | 30 | 3 |
| Measurements | 1 | 1 | 200 |  | 322 |
| Effective voxel size (mm) | 1.0x1.0x7.0 | 1.0x1.0x1.0 | 3.3x3.3x3.0 | 2.0x2.0x2.0 | 3.3x3.3x3.0 |
| Acquisition time (TA) | 0.25 | 4:26 | 6:44 | 2:14 | 1:00 |

#### ACR phantom

The ACR phantom is a commonly used phantom for QA. It uses a standardized imaging protocol with standardized MRI parameters (for an overview, see (ACR, 2005, 2008)). The protocol tests geometric accuracy, high-contrast spatial resolution, slice thickness accuracy, slice position accuracy, image intensity uniformity, percent-signal ghosting and low-contrast object detectability.

#### Phantom holder

At the beginning, both phantoms were manually aligned in the scanner and fixated using soft foam rubber pads. The alignment of the phantoms was evaluated by the radiographer performing the measurement and – if necessary – corrected using the localizer scan. To reduce spatial variance related to different placements of the phantom in the scanner and to decrease the time-consuming alignment procedure, we developed a styrofoam^TM^ phantom holder (Fig. 7). The phantom holder allowed a more time-efficient and standardized alignment of the phantoms within the scanner on the one hand. The measurement volumes of subsequent MR sequences could be placed automatically in the center of the phantom. On the other hand, the variability of QA statistics, related to different phantom mountings, was strongly reduced. This allowed a more sensitive assessment of MRI scanner stability (see Fig. 8 left).

**Fig. 7:**
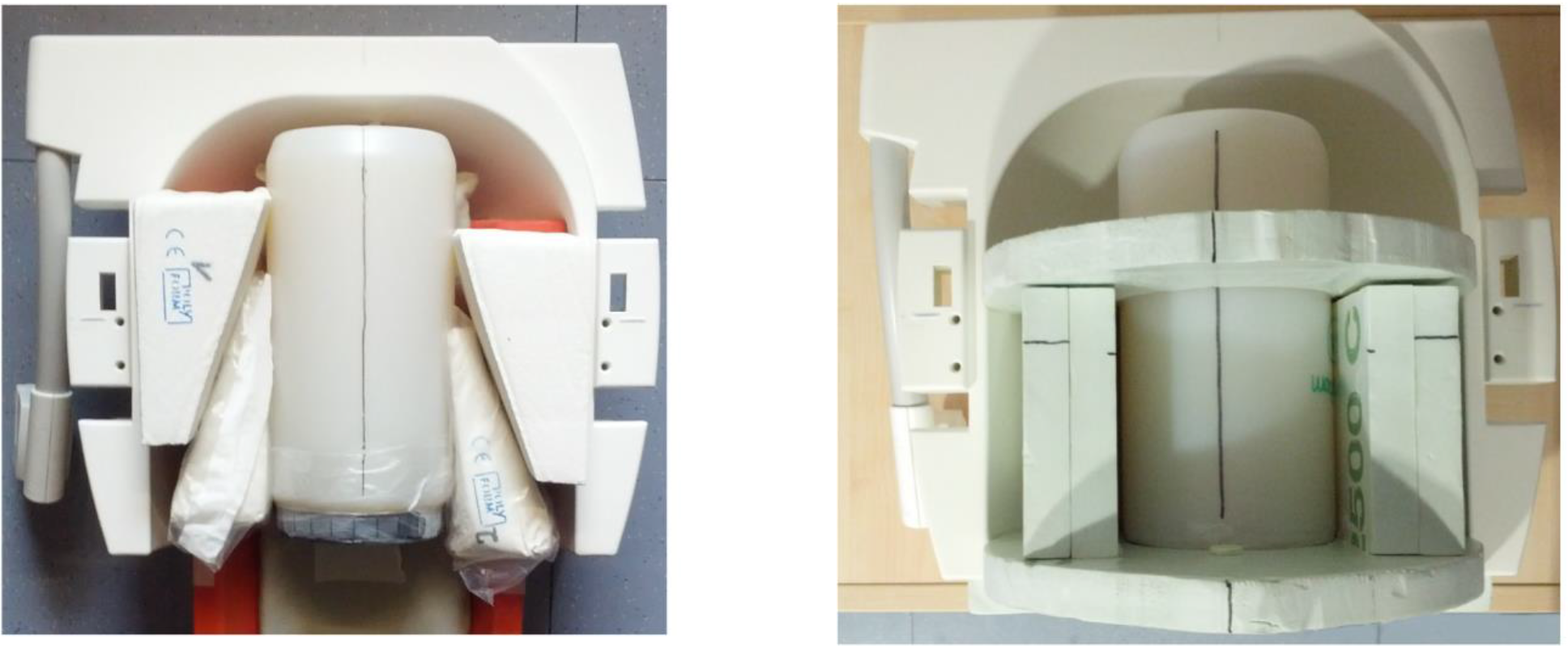
Manual alignment of the gel phantom using soft foam rubber pads (left) and more reliable alignment of the phantom using a Styrofoam holder (right).

**Fig. 8:**
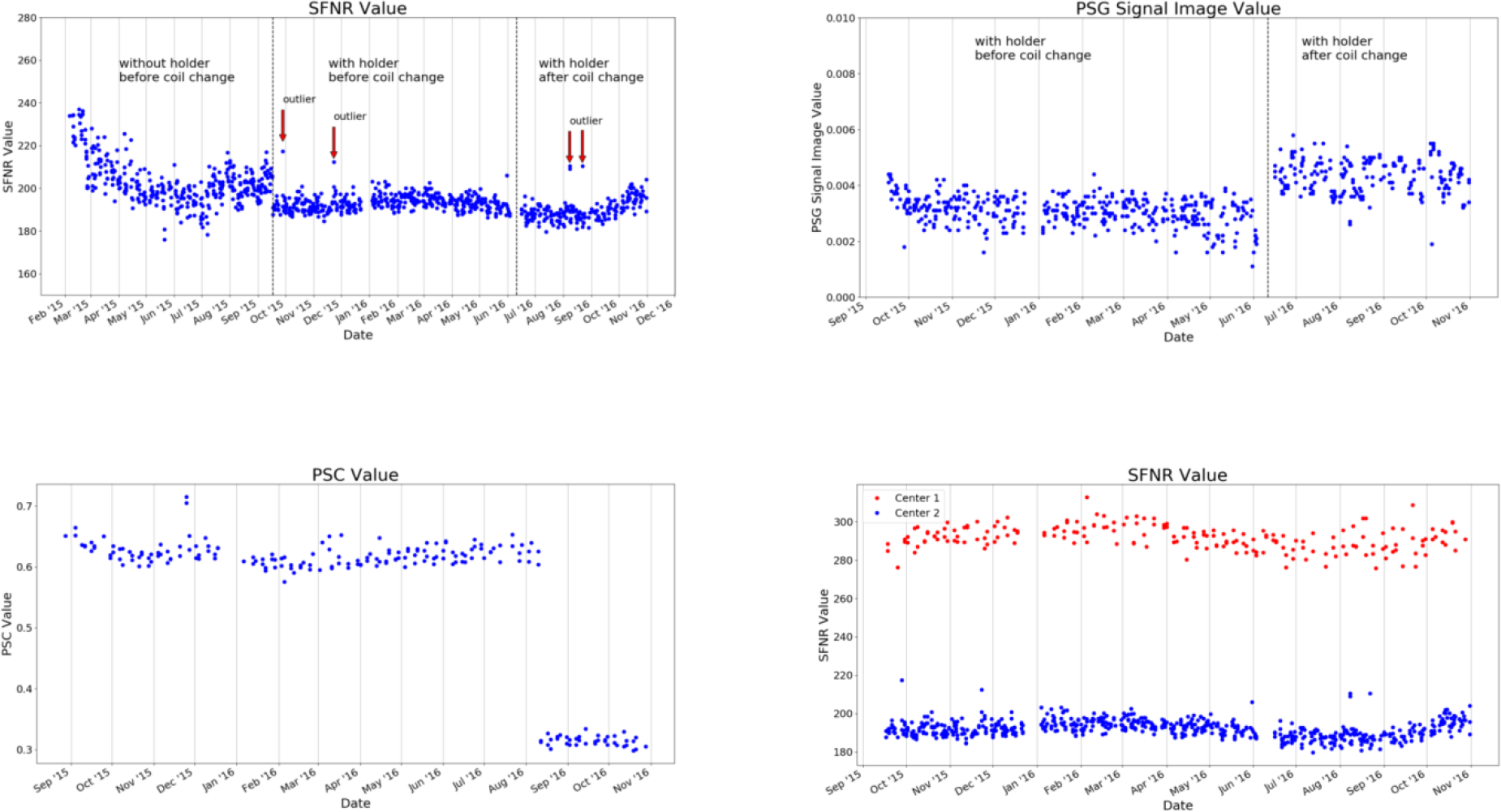
Selected QA statistics from gel phantom measurements. The data was collected over a duration of >1.5 years (Feb 2015 – Dec 2016) during the set-up of a large longitudinal imaging study (FOR 2107, Kircher et al. (2018)). Top left: After the implementation of a phantom holder (Oct 2015), the variance of many QA statistics was considerably reduced, as exemplarily shown for the signal-to-fluctuation-noise ratio (SFNR). This made it possible to detect outliers in future measurements (defined as 4 times SD of the mean; red arrows). Top right: In June 2016, the gradient coil had to be replaced. This had a major impact on the percent signal change (PSC). Bottom left: Changes in the MRI sequence, such as the introduction of the “prescan normalization” option that is used to make corrections for non-uniform receiver coil profiles prior to imaging, has a significant impact on the MRI data. This can be quantified using phantom data, as seen in the PSC. Bottom right: Imaging data in the study was collected at two different scanners. The scanner characteristics can also be determined using QA statistics, as shown for the SFNR. (For a detailed description of QA statistics, see Vogelbacher et al.(2018)).

In Fig. 8, we present selected QA data (from the gel phantom) collected over a duration of 22 months (Feb 2015 – Dec 2016) during the set-up of a large longitudinal imaging study (FOR 2107, Kircher et al. (2018)). The analysis of phantom data is able to show that changes in the QA-protocol (such as the introduction of a phantom holder, Fig. 8 top left), technical changes of a scanner (such as the replacement of the MRI gradient coil, Fig. 8 top right) or changes in certain sequence parameters (such as adding the prescan normalization option, Fig. 8 bottom left), impact many of the QA statistics in a variety of ways. It is also possible to use QA statistics to quantify the data quality of different MRI scanners (Fig. 8 bottom right). In summary, this exemplary selection of data shows the importance of QA analyses to assess the impact external events on the MRI data. The normal ranges of many QA statistics drastically change whenever hardware or software settings are changed at a scanner – both in mean and variance.

## 4 Discussion

In this article, we described a tool, *LAB–QA2GO*, for the fully-automatic quality assessment of MRI data. We developed two different types of QA analyses, a phantom and a human data QA pipeline. In its present implementation, *LAB–QA2GO* is able to run an automated QA analysis on data of ACR phantoms and gel phantoms. The ACR phantom is a widely used phantom for QA of MRI data. It tests in particular spatial properties, e.g. geometric accuracy, high-contrast spatial resolution or slice thickness accuracy. The gel phantom is mainly used to assess the temporal stability of the MRI data. For phantom data analysis, we used a wide array of previously described QA statistics (for an overview see e.g. Glover et al. (2012)). Although the main focus of the QA routines was the analysis of the phantom datasets, we additionally developed routines to analyze the quality of human datasets (without any pre-processing steps). *LAB–QA2GO* was developed in a modular fashion, making it easily possible to modify existing algorithms and to extend the QA analyses by adding self-designed routines. The tool is available for download on github^3^. License fees were avoided by using only open source software that was exempt from charges.

*LAB–QA2GO* is ready-to-use in about 10 minutes. Only a few configuration steps have to be performed. The tool does not need any further software or hardware requirements. *LAB– QA2GO* can receive MRI data either automatically (“network approach”) or manually (“stand-alone approach”). After sending data to the *LAB–QA2GO* tool, analysis of MRI data is performed automatically. All results are presented in an easy readable and easy-to-interpret web based format. The simple access via web-browser guarantees a user friendly usage without any specific IT knowledge as well as the minimalistic maintenance work of the tool. Results are presented both tabular and in graphical form. By inspecting the graphics on the overview page, the user is able to detect outliers easily. Potential outliers are highlighted by a warning sign. In each overview graph, an acceptance range (green area) is viable. This area can be defined for each graph individually (except for the ACR phantom because of the fixed acceptance values defined by the ACR protocol). To set up the acceptance range for a specific MRI scanner, we recommend some initial measurements to define the acceptance range. If a measurement is not in this range this might indicate performance problems of the MRI scanner.

Different QA protocols that assess MRI scanner stability are described in the literature, mostly designed for large-scale multicenter studies (for an overview see e.g. Glover et al. (2012)). Many of these protocols and the corresponding software tools are openly available. This allows in principle the flexible set-up of a QA protocol adapted for specific studies. The installation of these routines, however, is often not easy. The installation therefore often requires a fair level of technical experience, e.g. to install additional image processing software or to deal with specific software versions or hardware requirements. *LAB–QA2GO* was therefore developed with the aim to create an easily applicable QA tool. It provides on the one hand a fully automated QA pipeline, but is on the other hand easy to install on most imaging systems. Therefore, we envision that the tool might be a tailor-made solution for users without a strong technical background or for MRI laboratories without support of large core-facilities. Moreover, it also gives experienced users a minimalistic tool to easily calculate QA statistics for specific studies.

We outlined several possible application scenarios for the *LAB–QA2GO* tool. It can be used to assess the quality of MRI data sets acquired in small (with regard to sample size and study duration) neuroimaging studies, to standardize MRI scanners in multicenter imaging studies or to assess the long-term performance of MRI scanners. We outlined the use of the tool presenting data from center-specific QA protocol. These data showed that it was possible to detect outliers (i.e. bad data quality at some time points), to standardize MRI scanner performance and to evaluate the impact of hardware and software adaptation (e.g. the installation of a new gradient coil).

In the long run, the successful implementation of a QA protocol for imaging data does not only comprise the assessment of MRI data quality. QA has to be implemented on many different levels. A comprehensive QA protocol also has to encompass technical issues (e.g. monitoring of the temporal stability of the MRI signal in particular after hardware and software upgrades, use of secure database infrastructure that can store, retrieve, and monitor all collected data, documentation of changes on the MRI environment for instance with regard to scanner hardware, software updates) and should optimize management procedures (e.g. the careful coordination and division of labor, the actual data management, the long-term monitoring of measurement procedures, the compliance with regulations on data anonymity, the standardization of MRI measurement procedures). It also has to deal, especially at the beginning of a study, with the study design (e.g. selection of functional MRI paradigms that yield robust and reliable activation, determination of the longitudinal reliability of the imaging measures). Nonetheless, the fully-automatic quality assessment of MRI data constitutes an important part of any QA protocol for neuroimaging data.

In a future version of the tool we will add more possibilities to locate the unique identifier in the data e.g. by the use of different DICOM header values. We also will work on the automatic detection of the MR scanning parameters to start the corresponding QA protocol.

With *LAB–QA2GO* we hope to provide an easy-to-use toolbox that is able to calculate QA statistics without high effort.

## Footnotes

1 Some QA algorithms require e.g. the installation of standard image processing tools (e.g. Artifact Detection Tool (http://web.mit.edu/swg/software.htm); PCP Quality Assessment Protocol (Zarrar et al., 2015)) while others are integrated in different imaging tools (Mindcontrol (https://github.com/akeshavan/mindcontrol);BXH/XCEDE (Gadde et al., 2012)). Some pipelines can be integrated to commercial programs, e.g. MATLAB (CANlab (https://canlab.github.io/); ARTRepair), or large image processing systems (e.g. XNat (Marcus DS, Olsen TR, Ramaratnam M et al., 2007); C-Mind) of which some had own QA routines. Other QA pipelines can only be used online, by registering with a user account and uploading data to a server (e.g. LONI (Petros Petrosyan, Samuel Hobel, Andrei Irimia, John Van Horn, 2016)). Commercial software tools (e.g. BrainVoyager (Goebel, 2012)) mostly have their own QA pipeline included. Also some Docker based QA pipeline tools exist (e.g. MRIQC (Esteban et al., 2017)).

2 Virtual machines are common tools to virtualize a full system. The hypervisor for a virtual machine allocates an own set of resources for each VM of the host pc. Therefore, each VM is fully isolated. Based on the isolated approach of the VM technology, each VM has to update its own guest operating system. Another virtualization approach could have been based on Linux containers (e.g. Docker). Docker is a computer program that performs operating-system-level virtualization. This hypervisor uses the same resources which were allocated for the host pc and isolates just the running processes. Therefore, Docker only has to update the software to update all containers. For our tool we wanted to have a fully isolated system. Fixed software versions independent of the host pc are more likely to guarantee the functionality of the tool.

3 Github: https://github.com/vogelbac

